# Parallel evolution of direct development in frogs – Skin and thyroid gland development in African Squeaker Frogs (Anura: Arthroleptidae: *Arthroleptis*)

**DOI:** 10.1101/2020.09.07.286476

**Authors:** Benjamin Naumann, Susan Schweiger, Jörg U. Hammel, Hendrik Müller

## Abstract

Cases of parallel evolution offer the possibility to identify adaptive traits and to uncover developmental constraints on the evolutionary trajectories of these traits. The independent evolution of direct development, from the ancestral biphasic life history in frogs is such a case of parallel evolution. In frogs, aquatic larvae (tadpoles) differ profoundly from their adult forms and exhibit a stunning diversity regarding their habitats, morphology and feeding behaviors. The transition from the tadpole to the adult is a climactic, thyroid hormone (TH)-dependent process of profound and fast morphological rearrangement called metamorphosis. One of the organ systems that experiences the most comprehensive metamorphic rearrangements is the skin. Direct-developing frogs lack a free-swimming tadpole and hatch from terrestrial eggs as fully formed froglets. In the few species examined, development is characterized by the condensed and transient formation of some tadpole-specific features and the early formation of adult-specific features during a “cryptic” metamorphosis. In this study we show that skin in direct-developing African squeaker frogs (*Arthroleptis*) is also repatterned from a tadpole-like to an adult-like histology during a cryptic metamorphosis. This repatterning correlates with an increase of thyroid gland activity. A comparison with data from the Puerto Rican coqui (*Eleutherodactylus coqui*) reveals that direct development might have evolved in parallel in these frogs by a comparable heterochronic shift of thyroid gland activity. This suggests that the development of many adult-features is still constrained by the ancestral dependency on thyroid hormone signaling.

## Introduction

How do developmental constraints influence phenotypic evolution? A promising way to approach this question is studying cases of parallel evolution (Schluter, Clifford, Nemethy, & McKinnon, 2004). Parallel evolution can be defined as the independent origin of similar (derived) traits in two or more taxa sharing a common ancestry and bauplan (Futuyma & Kirkpatrick, 2017; Simpson, 1961). Investigating patterns of parallel evolution allows to identify adaptive traits and to uncover developmental constraints on the evolutionary trajectories of these traits (Schluter et al., 2004). A potential case of parallel evolution is the repeated, independent origin of direct development within frogs and other amphibians (San Mauro et al., 2014; Schweiger, Naumann, Larson, Möckel, & Müller, 2017; Thibaudeau & Altig, 1999; Wake & Hanken, 1996). Direct development is characterized by the loss of an aquatic larval phase (Hall & Olson, 2003) and may have evolved in response to environmental conditions that restricted the availability of suitable habitats for aquatic development (Goin & Goin, 1962; Liedtke et al., 2017; Müller et al., 2013; ten Brink, Onstein, & de Roos, 2020).

In frogs, direct development evolved several times independently (Figure 1A), making them a suitable model system to unravel developmental constraints that underlay patterns of parallel evolution (Goldberg, Candioti, & Akmentins, 2012; Heinicke et al., 2009). The ancestral biphasic life history of frogs includes a free-swimming larval stage called tadpole (McDiarmid & Altig, 1999). In contrast to other amphibian larvae, tadpoles differ extremely from their adult forms and exhibit a stunning diversity regarding their habitats, morphology and feeding behaviors (Altig & McDiarmid, 1999). In biphasic anurans, embryonic development from a fertilized egg to a tadpole happens more or less gradually, similar to other vertebrate groups. In contrast, the transition from the tadpole to the adult is a climactic, thyroid hormone (TH)-dependent process of profound and fast morphological rearrangement called metamorphosis (Shi, 1999) (Figure 1B). In direct developing species, development from the embryo to the adult frog appears more gradual without an obvious climactic phase (Callery, Fang, & Elinson, 2001) (Figure 1B). Tadpole-specific traits such as the larval skeleton and its associated musculature, the cement glands, the lateral line system and the coiled intestine are nearly or completely reduced (Callery et al., 2001; Hanken, Klymkowsky, Alley, & Jennings, 1997). However, morphological studies on the embryonic development of direct developing frogs are limited to only a few species (Goldberg et al., 2012; Goldberg, Taucce, Quinzio, Haddad, & Candioti, 2020; Hanken, Klymkowsky, Summers, Seufert, & Ingebrigtsen, 1992; Kerney, Meegaskumbura, Manamendra□Arachchi, & Hanken, 2007; Schweiger et al., 2017). The only species in which direct development has been investigated in greater detail is the Puerto Rican coqui, *Eleutherodactylus coqui* Thomas, 1966 (thyroid gland: Laslo, Denver, & Hanken, 2019; Callery & Elinson, 2000; Elinson & Fang, 1998; Jennings & Hanken, 1998; Fang & Elinson, 1996; limb development: Hanken et al., 2001; cranial development: Hanken et al., 1997; Kerney, Gross, & Hanken, 2010; Olsson, Moury, Carl, Håstad, & Hanken, 2002; Schlosser, Kintner, & Northcutt, 1999; Schlosser & Roth, 1997; gross embryonic development: Townsend & Stewart, 1985).

**Figure 1.**
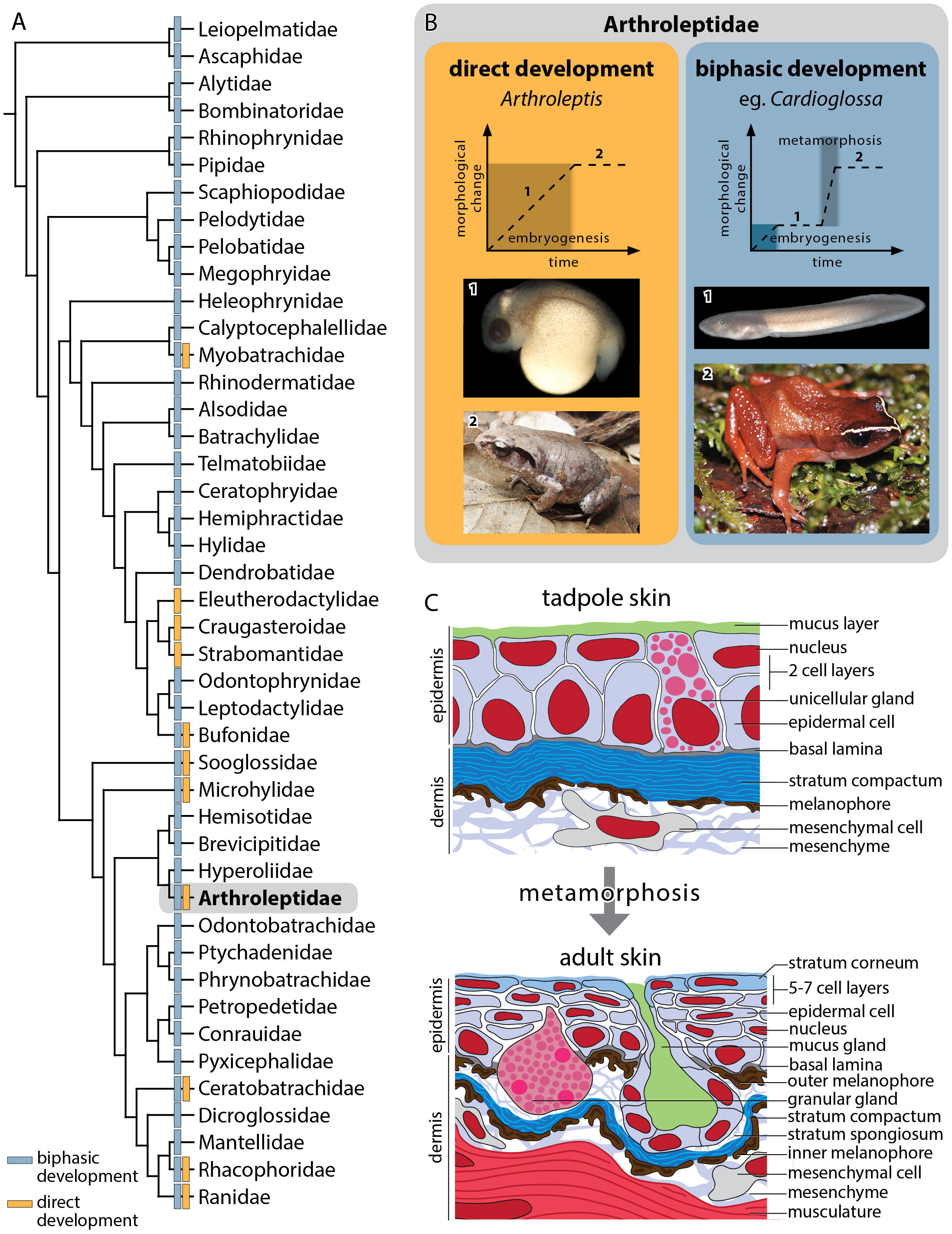
A, a family-level phylogeny of recent frogs based on Feng et al., 2017 (Feng et al., 2017). Blue boxes indicate the presence of an ancestral biphasic live history while orange boxes indicate direct development. B, a schematic graph showing the gradual morphological change during ontogeny of the direct developing *Arthroleptis* (*A. wahlbergii*; 1, embryo; 2, adult frog) and the more climactic metamorphic change in its biphasic sister genus *Cardioglossa* (*C. manengouba*; 1, tadpole; 2, adult frog). Photographs are not to scale. C, schematic diagrams of the organization of the tadpole and adult frog skin.

Although direct-developing frogs have departed profoundly from their ancestral ontogeny (Hanken et al., 1997; Schlosser, 2008; Schweiger et al., 2017), studies in *E. coqui* have shown that several developmental events are still under the control of TH (Callery & Elinson, 2000; Hanken & Summers, 1988) and that it undergoes a cryptic metamorphosis (Callery & Elinson, 2000; Ziermann & Diogo, 2014). The ontogenetic repatterning seen in direct-developing frogs is thought to be the result of changes in the expression of TH, which leads to changes in timing of TH-dependent events during development (“heterochronic shift” hypothesis). At least some developmental events are thought to have become decoupled from TH-regulation (“loss of constraint” hypothesis), but so far data are only available for *E. coqui* and the general influence of TH during the ontogeny of other direct-developing frogs remains unclear.

An interesting system to study putative developmental constraints on the evolution of direct development is the skin. The frog skin hosts a complex microbiome and plays major roles in immune response, pathogen defense, respiration, osmoregulation, camouflage and even reproduction (Douglas, Hug, & Katzenback, 2020; Fernandes et al., 2011; Huang et al., 2016; Katz, 1986; Varga, Bui-Marinos, & Katzenback, 2019). Among all tissues that remodel during metamorphosis the skin exhibits the most extreme changes regarding histology and gene expression (Yoshizato, 1992). It is exposed to completely different environments (aquatic vs. terrestrial) depending on life history phases and often exhibits tadpole- and adult-specific adaptations (Quinzio & Goldberg, 2019). The skin of pre-metamorphic tadpoles is a mostly two-layered epithelium with unicellular glands and an outer mucus layer (Duellman & Trueb, 1994). The underlying dermis consists of a stratum compactum and some melanophores in the mesenchyme beneath (Figure 1C). During pro-metamorphosis, the cells of the epidermis degenerate except for cells attached to the basal lamina. These basal cells start to proliferate during metamorphic climax and build up the post-metamorphic five- to seven-layered adult frog epidermis with an outer keratinized cell layer (stratum corneum) and different multicellular gland types. The underlying dermis contains an outer (exterior to the stratum compactum) and an inner (interior to the stratum compactum) melanophore layer (Figure 1C) (Fox, 1985; Kinoshita & Sasaki, 1994; Tamakoshi, Oofusa, & Yoshizato, 1998). Metamorphic changes in biphasic species are initiated by thyroid hormone (TH) production (Brown & Cai, 2007; Kulkarni, Singamsetty, & Buchholz, 2010; Tata, 2006). Thyroid hormone is also involved in several developmental processes in *E. coqui* indicating the presence of a cryptic metamorphosis in direct developing species (Callery & Elinson, 2000; Hanken et al., 1992; Kulkarni et al., 2010; Schlosser, 2008).

In this study, we provide detailed data on skin development in direct developing African Squeaker frogs (*Arthroleptis*) using histological and immuno-histochemical techniques. To evaluate if potential changes in the skin and thyroid gland histology of *Arthroleptis* are due to direct development or instead shared by biphasic Arthroleptidae, we investigated the same tissues in a *Cardioglossa* tadpole, the sister genus of *Arthroleptis*. Additionally, we describe thyroid gland development in *Arthroleptis* and infer its activity based on morphology and morphometric measurements.

## Results

Many aspects of the development of the embryonic pigmentation pattern, skin and thyroid glands are similar in *A. wahlbergii* and *A. xenodactyloides*. Differences between the two species are mentioned where present.

### Development of the pigmentation pattern in *Arthroleptis* embryos

Embryos until mid TS5 lack any obvious, externally visible pigmentation and have a white-yellowish color (Figure 2A; see also Schweiger et al., 2017). However, very few melanophores were found in the epidermal layer in histological sections of an embryo at TS4 (see Figure 3A4). At late TS5/early TS6 melanophores are recognizable extending dorso-ventrad until the ventral border of the eye in the head region and until the dorsal-most quarter of the lateral body wall in the trunk region (Figure 2B). Melanophores at this stage are spindle-shaped with long, thin extensions (Figure 2B’). From late TS7 to TS8, melanophores continue to extend ventrad in the head and the trunk regions (Figure 2C, D) until they completely cover the lateral body wall from TS9 on (Figure 2E, F). At this stage, the body wall has completely enclosed the yolk (Figure 2E3). At TS11, the density of melanocytes of the head region starts to increase resulting in a darkening of the skin (Figure 2G). Additionally, another type of melanophores becomes visible all over the body. These melanophores are smaller, with a more spherical shape and with only a few short or no extensions (Figure 2G’). At TS12, melanocyte density continues to increase dorso-ventrad until the whole lateral body wall is heavily pigmented at TS15 (Figure 2H-K).

**Figure 2.**
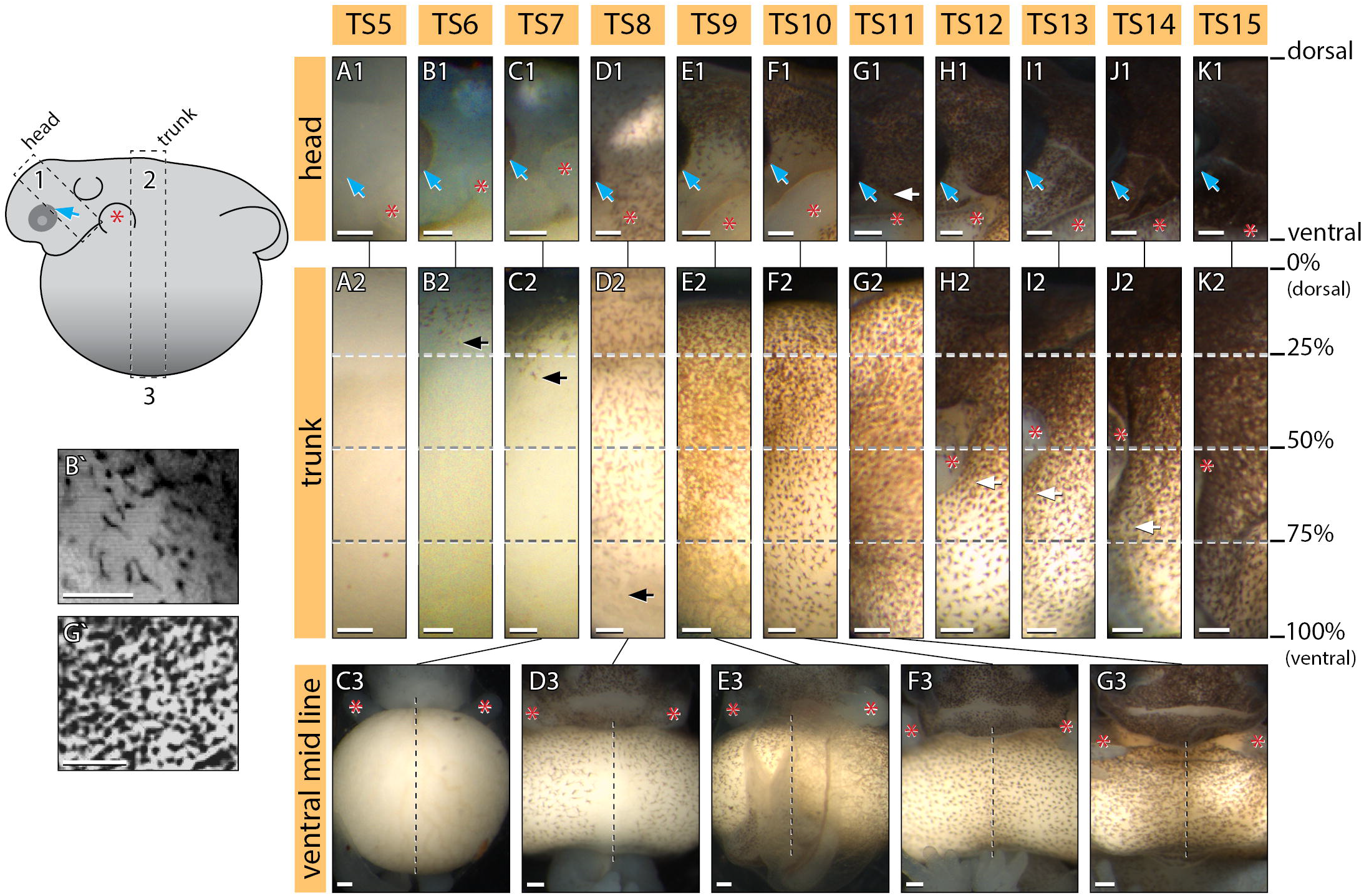
A-G, Photographs of the skin pigmentation of different developmental stages of *Arthroleptis wahlbergii*. The schematic embryo on the left illustrates the photographed regions (1, 2 and 3). Blue arrows indicate the position of the eye, red asterisks the position of the forelimb. The dotted grey lines and percentage number indicate the distance between the back (dorsal, 0%) and ventral midline (ventral, 100%). The black arrow in A2-D2 indicates the migration distance of the first melanophore type (shown in B’). The white arrow in H2-J2 indicates the migration distance of the second melanophore type (shown in G’). The black dotted line in C3-G3 indicates the ventral midline. Scale bar is 200 μm.

**Figure 3.**
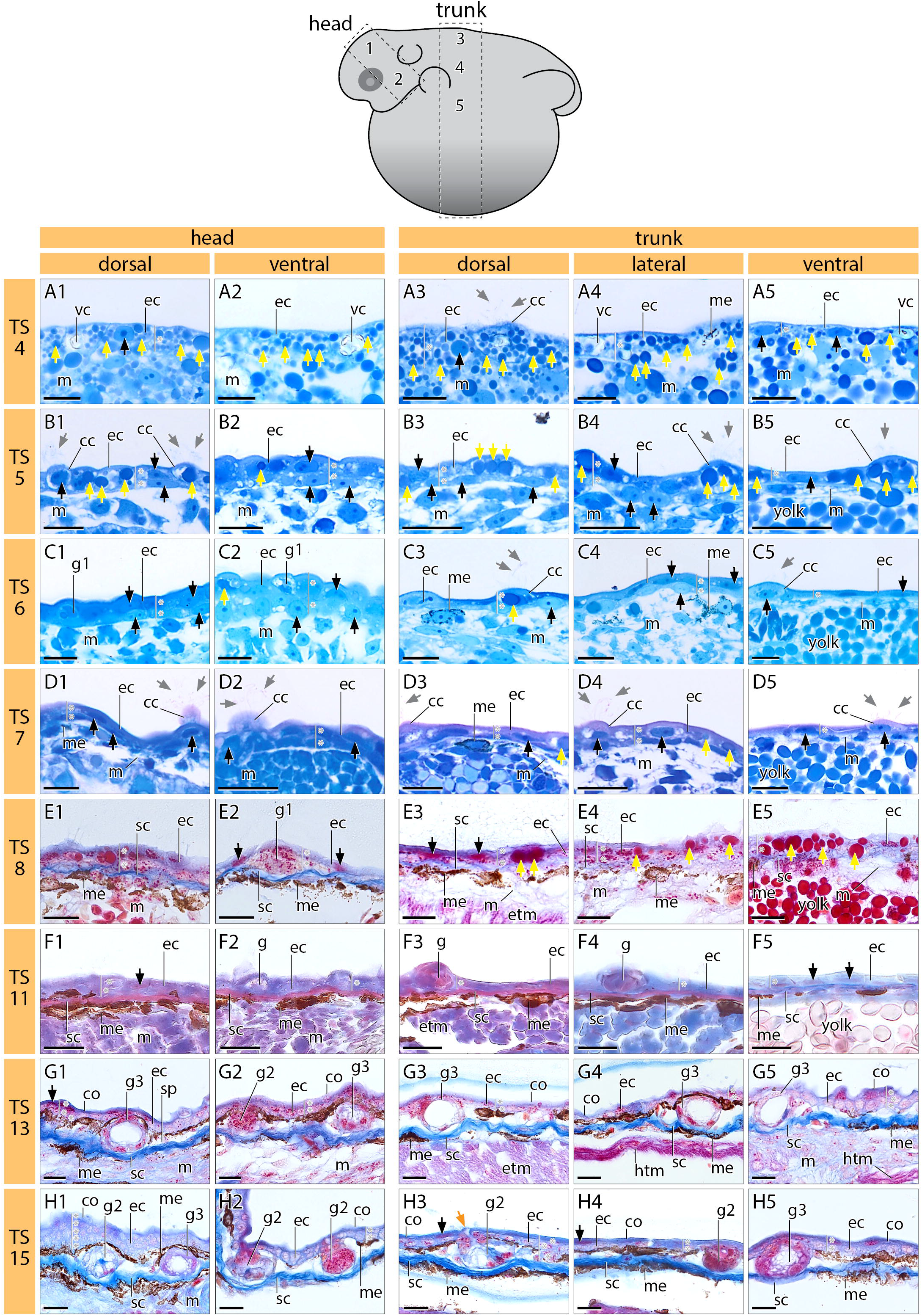
Histological cross sections of the skin of *Arthroleptis wahlbergii* at different developmental stages. The schematic embryo on top illustrates the photographed regions (1-5). Grey arrows indicate the epidermal cilia. Yellow arrows indicate intracellular yolk droplets. Black arrows indicate epidermal nuclei. The yellow arrow in H3 indicates the glandular duct. The number of epidermal cellular layers is indicated by a grey line and asterisks. Scale bar is 20 μm. cc, ciliated cell; co, stratum corneum; ec, epithel cell; etm, epaxonic trunk muscles; g, multicellular gland progenitor; g1, unicellular gland; g2, multicellular granular gland; g3, multicellular mucus gland; htm, hypaxonic trunk muscles; m, mesenchyme; me, melanophore; sc, stratum compactum; sp, stratum spongiosum; vc, vacuole.

In summary, pigmentation development is characterized by a first dorso-ventrad wave of melanophore development starting at late TS5 and a second wave made up by a morphologically different type of melanophores starting at TS11.

### Skin development in *Arthroleptis*

TS4 (Figure 3A) – The epidermis is single-layered with a sometimes indistinct border between the underlying cells and tissues. In the dorsal body region, cells are large and cuboid with spherical nuclei. Ventrally, cells are flattened and nuclei are ovoid to spindle-shaped. The cells possess high amounts of intracellular yolk granules and some seem to have large vacuoles with dark pigments. Ciliated cells are found in a scattered pattern all over the embryo. Very few melanophores can be found intercalating between epidermal cells. A dense, undifferentiated mesenchyme lies beneath the epidermis.

TS5 (Figure 3B) – The dorsal epidermis of the head as well as of the dorso-lateral trunk is two-layered while the ventral epidermis of the trunk, covering the yolk sac, is single-layered. Most cells are slightly flattened with ovoid nuclei. The amount of intracellular yolk granules has decreased compared to TS4 but is still high. The number of ciliated cells has increased compared to TS4. A sharp border between epidermal cells and the underlying mesenchyme is recognizable.

TS6 and TS7 (Figure 3C, D) – The number of epidermal layers has not changed compared to TS5. Some unicellular mucus glands are recognizable. The amount of intracellular yolk granules has further decreased compared to previous stages. A higher number of melanophores with long cytoplasmic extensions are present in the mesenchyme directly beneath the epidermis.

TS8 (Figure 3E) – The number of epithelial layers is similar to previous stages. Epidermal cells in both layers are flattened and show spindle-shaped nuclei. Some hypertrophied cells with a granular content that might represent unicellular glands are present in the skin of the head. Intracellular yolk granules are only detectable in the trunk epidermis. In the head, a distinct stratum compactum is present beneath the epidermis with a dense layer of inner melanophores interior to it. In the trunk, the stratum compactum is most distinct in the dorsal region and becomes gradually less obvious in lateral and ventral areas. This pattern is mirrored by the density of associated inner melanophores.

TS11 (Figure 3F) – The epidermis has become single-layered again in the head and dorsolateral trunk region and epidermal cells exhibit extremely flattened nuclei. In some scattered areas of the dorsal head, the epidermis is still two-layered. Intracellular yolk granules and ciliated cells are completely absent. Multicellular gland primordia, consisting of several compact cells, can be found all over the body. In the head, a few outer melanocytes are present between the stratum compactum and basal epidermal cells. In the trunk, the stratum compactum appears denser compared to the previous stage and the density of inner melanophores has increased. It is still lowest in the ventral trunk region.

TS13 (Figure 3G) – The skin exhibits many features of an adult frog skin. The epidermis is two - to three-layered. The majority of the basal epidermal cells are cuboid. The apical cell layer is flattened and is stained dark-blue in Azan-stained section, indicating an increased keratinization and the presence of a stratum corneum. The stratum corneum is more obvious in the head compared to the trunk epidermis. Multicellular mucus and granular glands can be found in various regions of the body. They are localized within a thin stratum spongiosum beneath the basal epidermal cell layer. Mucus glands are outlined by a single cell layer with an inner lumen. Granular glands are recognizable as large sacs filled with many granulated cells. The density of outer melanophores within the now developed stratum spongiosum has increased compared to previous stages.

TS15 (Figure 3H) – Hatching occurs at this stage and the skin of the froglet is similar to the skin of an adult *Arthroleptis* (Supplementary material 2). The thickness of the epidermis varies from two - to five cell layers but is three-layered in most areas. In the trunk region, the dark blue-stained stratum corneum is now also recognizable as a distinct cellular layer. Multicellular glands in the whole skin are more numerous and melanophore density has increased in the ventral trunk region compared to TS13.

In summary, the embryonic ectoderm and underlying mesenchyme at TS4 differentiated into a skin consisting of an epidermis and an underlying dermis both exhibiting pre-metamorphic tadpole-typical features at TS8. At TS11 the apical epidermal layers seem to degenerate to a certain degree while multicellular gland progenitors appear all over the body. The skin of embryos at TS13 and TS15 exhibit many features typical for the post-metamorphic frog skin.

### PCNA expression during epidermal development in *Arthroleptis*

Metamorphic skin remodeling is based on the degeneration of apical epidermal cells and the proliferation of remaining basal cells building the adult epidermis (Yoshizato, 1992). We therefore investigated the epidermal proliferation pattern in embryonic stages of *A. wahlbergii* using two different antibodies against proliferating cell nuclear antigen (PCNA). Signals from both antibodies were detected in the same tissues of two adjacent serial sections verifying antibody specificity. However, the signal from PCNA-1 antibody was always stronger than from the PCNA-2 antibody.

At TS8, scattered signals of the PCNA-1 antibody are detectable in nuclei of both, epidermal cell layers as well as some mesenchymal cells (Figure 4A, A’). Only a few PCNA-2 positive cells are detectable at this stage (Figure 4B, B’). At TS11, PCNA-1 and PCNA-2 signals are detectable in the majority of epidermal cells, indicating a strong increase in the proliferation rate of the epidermis (Figure 4C, D). A few PCNA-1 and PCNA-2 positive mesenchymal cells are also present. No antibody signals are detectable in the epidermis at TS13 and TS15, indicating a low proliferation rate (Figure 4E-G). In some multicellular glands however, PCNA-1 positive cells can be detected (Figure 4G).

**Figure 4.**
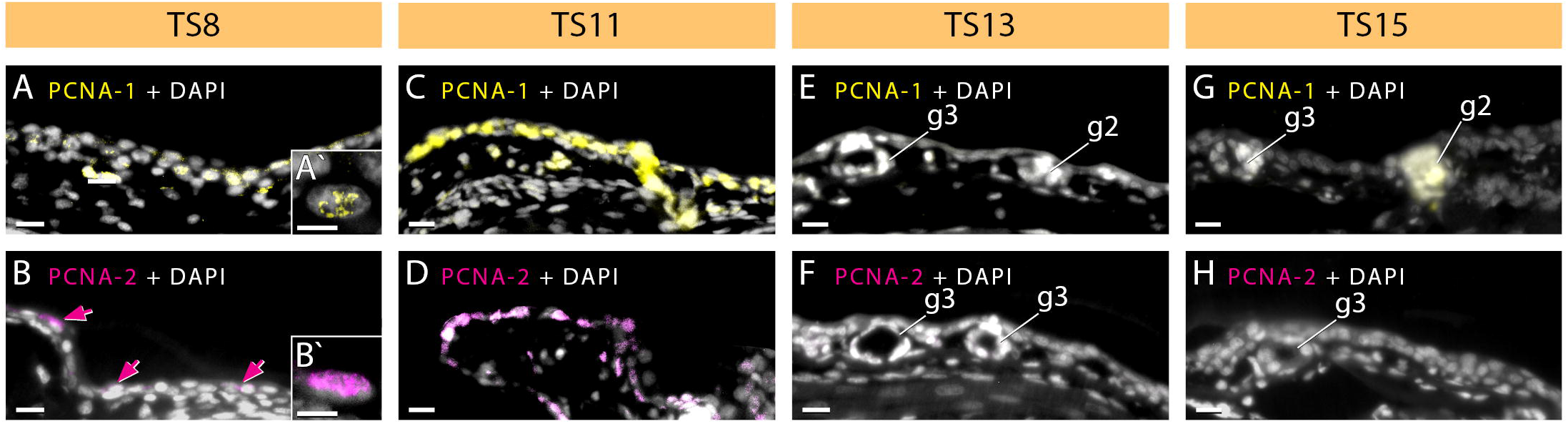
Fluorescent PCNA antibody staining in cross sections through the skin of *Arthroleptis wahlbergii* embryos at different developmental stages. Two different antibodies against PCNA have been used. PCNA-1 is colored in yellow and shown in A, C, E and G. PCNA-2 is colored in magenta and shown in B, D, F and H. Nuclei are stained with DAPI and colored in grey. A’ and B’ Show a close up of the PCNA signal within the nucleus. Magenta arrows in B indicate the scattered antibody signal. A-H, scale bar is 20 μm. A’ and B’, scale bar is 10 μm. g2, multicellular granular gland; g3, multicellular mucus gland.

In summary, PCNA reactivity (and therefore cell proliferation rate) of the skin is detectable in a scattered pattern at TS8, has increased tremendously at TS11 and is low or absent at TS 13 and TS15.

### Thyroid gland development in *Arthroleptis* embryos

The thyroid glands are paired, ovoid structures located at the lateral edges of the hyoid plate (Figure 5A-C). Primordia are first detectable at TS8. They appear as spherical, condensed cell masses including scattered lumina which lack an epithelial lining. The primordia are located on the left and right side ventrally to the lateral edges of the developing hyoid plate (Figure 5A, D). First developing follicles are recognizable at TS10/11. The cells of the follicular epithelial are cuboid with little cytoplasm and large spherical nuclei (Figure 5E). At later TS11, thyroid glands have elongated, adopting a mature, adult-like morphology (Figure 5B). The follicles have increased in size and number and the follicular epithelium is now organized in a columnar-pattern. Nuclei of follicular epithelial cells are spherical to ovoid-shaped. Some colloid, recognizable as a bluish substance in Azan-stained sections, is present within follicular lumina (Figure 5F). At TS13 the follicles have further increased in size and number. The cells of the follicular epithelium appear slightly flattened compared to the previous stage. No colloid is detectable in the investigated specimen (Figure 5G). At TS15 the thyroid glands have continued to elongate and are almost spindle-shaped (Figure 5C). The size and number of follicles have further increased and the follicular lumina are filled with colloid in the investigated specimen. Erythrocytes are recognizable around and in between follicles indicating a continuous vascularization of the thyroid gland (Figure 5H).

**Figure 5.**
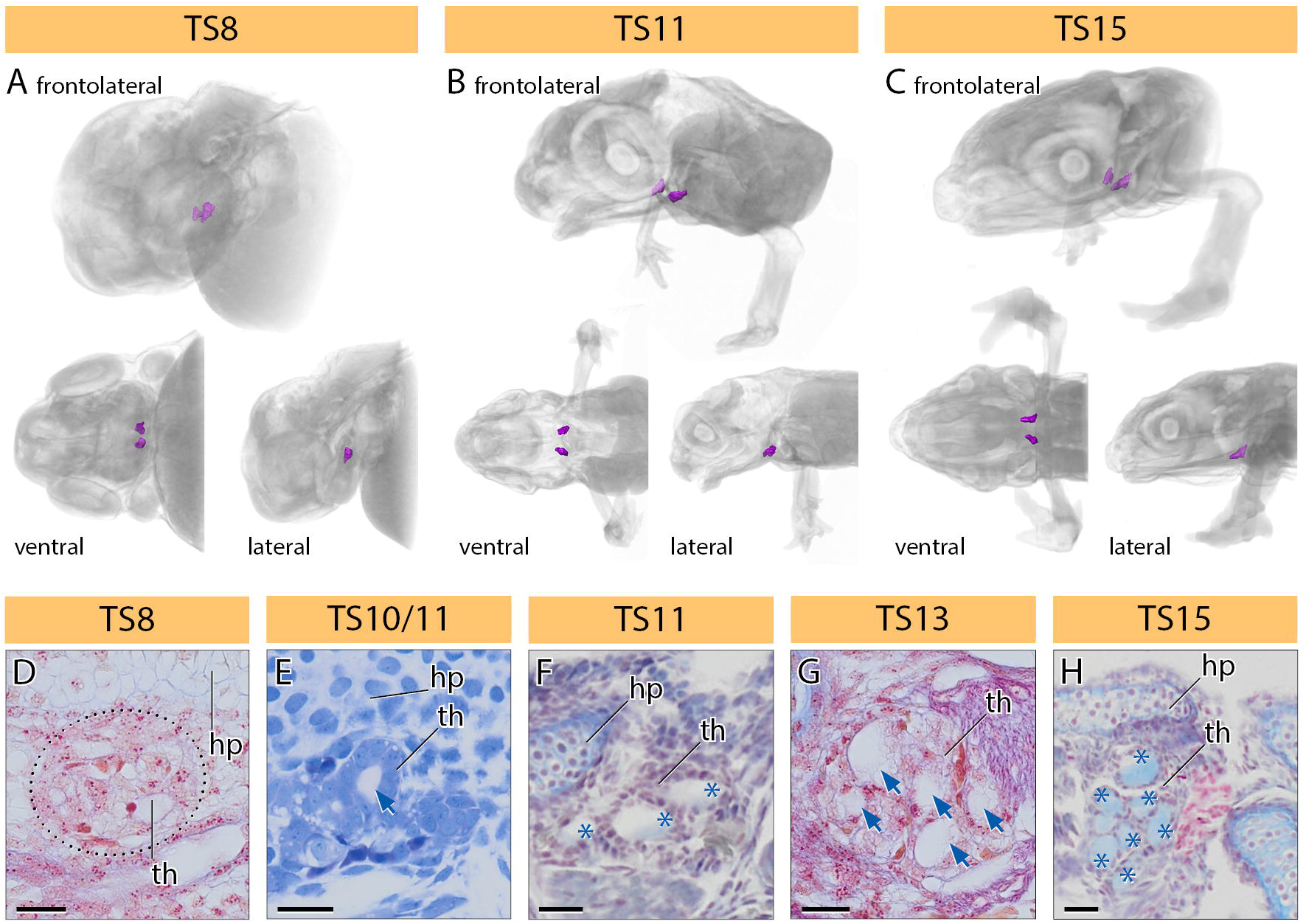
A-C volume renderings of μCT scans of different developmental stages of *Arthroleptis xenodactyloides*. The thyroid glands are colored in lilac. Renderings are not to scale. D-H, histological cross-sections of the ventral head region of *A. wahlbergii* embryos at different developmental stages. Blue arrows indicate absence of colloid within follicles. Blue asterisks indicate the presence of colloid within follicles. Scale bar is 25 μm. hp, hyoid plate; th, thyroid gland.

We therefore measured the cell height of the follicular epithelium as well as the number and diameter of follicles (Figure 6B-D). All measurements were taken from sections from the central region of the thyroid gland (Cruz & Fabrezi, 2020). In *A. wahlbergi* we observed a huge increase in the follicle cell height from TS8 to TS10/11, followed by a gradual decrease in TS 13 and TS15. In the two examined stages of *A. xenodactyloides* cells were smaller compared to *A. wahlbergii*. However, a similar but lower decrease in cell height from TS11 to TS15 is also recognizable (Figure 6B). Follicle numbers and follicle number are more or less similar in *A. wahlbergii* and *A. xenodactyloides* (Figure 6C, D). In *A. wahlbergi*, follicle number and diameter decrease from TS8 to TS10/11 and then increases gradually from TS10/11 to TS15. In the two specimens of *A. xenodactyloides*, an increase in follicle number and diameter can also be observed.

**Figure 6.**
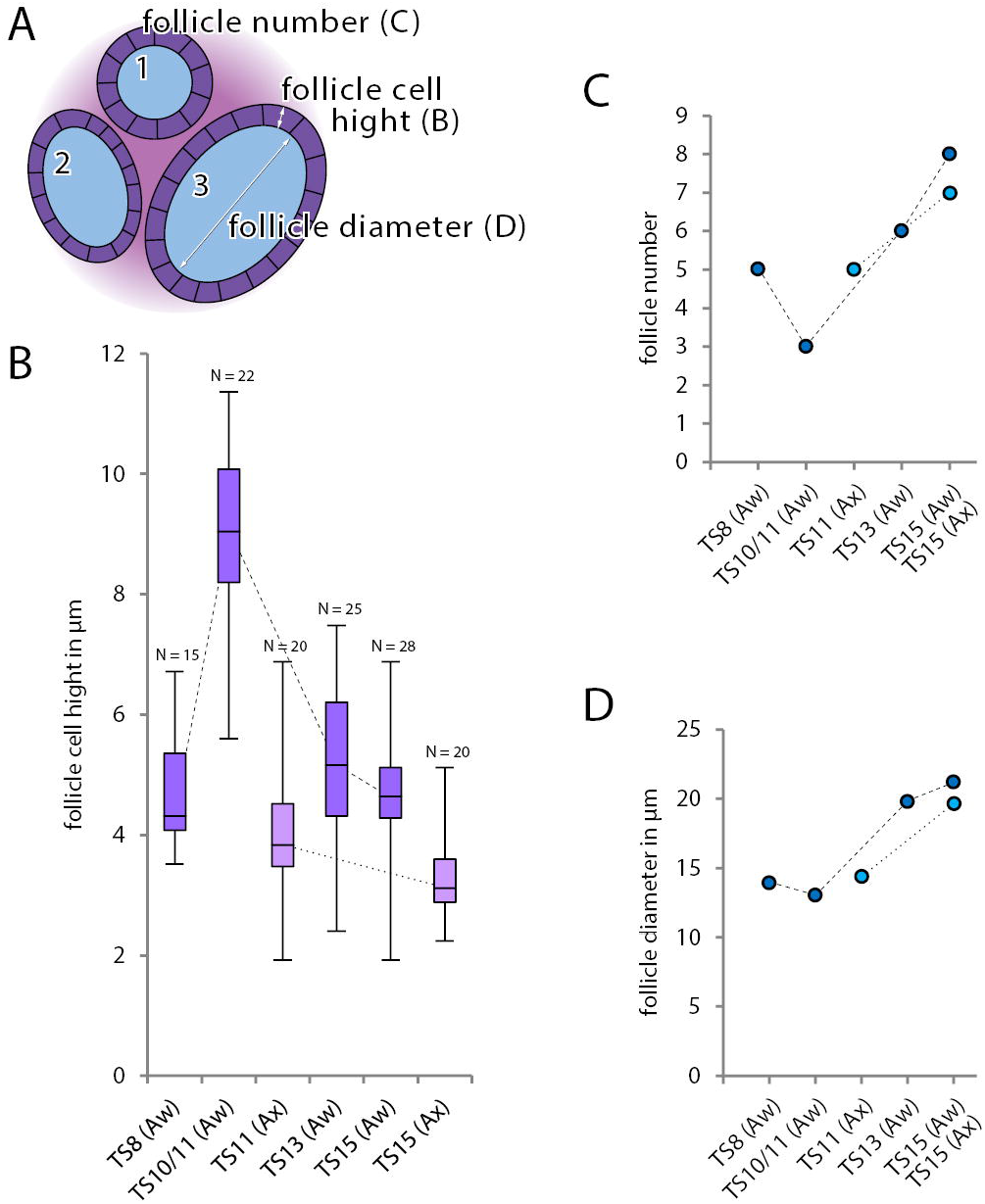
A, schematic illustration of thyroid gland follicels and the measured parameters. B, Box plot of the follicle cell height in *Arthroleptis* at different developmental stages. C, diagram of the follicle number in *Arthroleptis* at different developmental stages. D, diagram of the average follicle diameter in *Arthroleptis* at different developmental stages. Aw, *A. wahlbergii*; Ax, *A. xenodactyloides*.

In summary, our data indicate that thyroid glands differentiate histologically between TS8 and TS10/11, become highly active at TS11 (follicle cell height, first detected colloid) and develop and adult-like morphology between TS 13 and TS15 (increasing follicle diameter and number, decreasing cell height).

### Skin and thyroid gland histology in a *Cardioglossa* tadpole

The skin of *Cardioglossa* sp. at Gosner-stage 27 (Gosner, 1960) exhibits a pattern typical for the majority of tadpoles (McDiarmid & Altig, 1999). The epidermis is two-layered. The basal layer consists of cuboid cells with spherical nuclei while the apical layer is slightly flattened in the head but not in the trunk region (Figure 7A1-A5). A dark-blue stained extracellular layer resembling the keratinized apical cell layer present in late *Arthroleptis* embryos can be found. Some unicellular gland cells are present in between basal and apical cells (Figure 7A1, A3). A thick stratum compactum is present beneath the basal epidermal layer. Melanophores are associated with the stratum compactum. Their density decreases dorso-ventrad in the head as well as the trunk region (Figure 7A1-A5).

**Figure 7.**
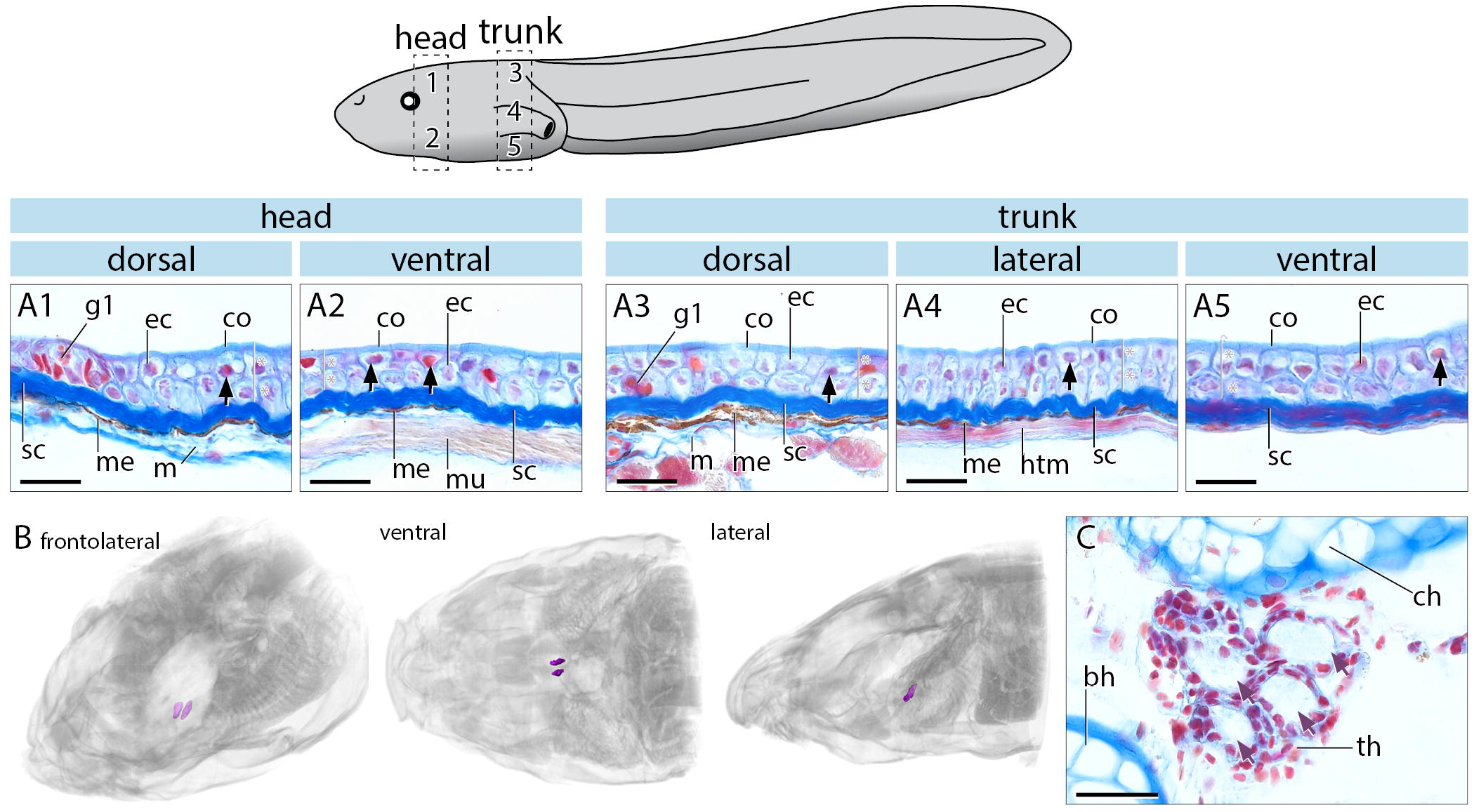
A, histological cross sections of the skin of a *Cardioglossa manengouba* tadpole at Gosner-stage 27. The schematic tadpole on top illustrates the photographed regions (1-5). Black arrows indicate epidermal nuclei. The number of epidermal cellular layers is indicated by a grey line and asterisks. B, volume renderings of a synchrotron scan of a *C. manengouba* tadpole at Gosner-stage 27. The thyroid glands are colored in lilac. Renderings are not to scale. C, histological cross sections of the ventral head region of the same tadpole as in A. Lilac arrows indicate absence of colloid within follicles. Scale bar is 20 μm. bh, basihyal; ch, ceratohyal; co, keratinized layer; ec, epithel cell; g1, unicellular gland; htm, hypaxonic trunk muscles; m, mesenchyme; me, melanophore; mu, muscle; sc, stratum compactum; th, thyroid gland.

The thyroid glands exhibit a more elongated olive-shape and are localized in the same position as described for *Arthroleptis* embryos (Figure 7B). At this developmental stage, well developed follicle outlines by slightly flattened epithelial cells can be recognized. No colloid or vascularization has been detected in the examined specimen. The flattened follicular epithelial cells and the absence of colloid and vascularization indicate a low activity of TH-production.

## Discussion

### *Arthroleptis* exhibits accelerated body wall fusion and a phase of metamorphic skin pigmentation repatterning that correlates with increased thyroid gland activity

The skin of frogs harbors a variety of chromatophores (Duellman & Trueb, 1994). In this study however we focus on the pattern of melanophores only. Different types of melanophores are present in the adult frog skin (Yasutomi, 1987). This adult pigmentation pattern is established via metamorphic changes in the dermal pigmentation, chromophore morphology, and biochemistry of the tadpole skin (Yasutomi, 1987). This indicates a correlation of TH production and metamorphic skin pigmentation repatterning in biphasic anurans (Smith-Gill & Carver, 1981).

In direct developing species, pigmentation patterning has only been described in *E. coqui* (Townsend and Stewart, 1985). At TS7, first melanophores appear in the trunk region and pigmentation increases slowly until it becomes heavy at TS10. Pigmentation of the head is delayed becoming heavy at TS12. This general process of melanophore patterning is also seen in *Arthroleptis* (Schweiger et al., 2017; this study) and we here provide the first detailed data on the morphology of different melanophore types during skin development in a direct developing frog species. In *Arthroleptis*, the first type of melanophore is spindle-shaped with long thin extensions. This type is typical of embryonic melanophores in biphasic species (Smith-Gill & Carver, 1981; Yasutomi, 1987). A second type of melanophore with a smaller, more circular-shape with a few short or no extensions appears at around TS11. This melanophore type resembles melanophores appearing during TH-mediated metamorphosis in biphasic species (Smith-Gill & Carver, 1981; Yasutomi, 1987). It is generally accepted that follicular cell height, follicle diameter and number (see Figure 6A) roughly reflect thyroid gland activity and TH production (Coleman, Evennett, & Dodd, 1968; Grim et al., 2009). Therefore, the appearance of this second melanophore type correlates with an increased thyroid activity. The timing of pigmentation repatterning and melanophore type appearance in *Arthroleptis* is very similar to the TH-mediated metamorphic transitions seen in the skin of biphasic species (Smith-Gill & Carver, 1981; Yasutomi, 1987).

Experimental studies on the pigmentation pattern (Elinson & Fang, 1998) and histological investigation of thyroid gland activity (Jennings & Hanken, 1998) in *E. coqui* led to the suggestion that TH is also involved in the closure of the pigmented body surface (Callery & Elinson, 2000; Elinson & Fang, 1998). While the overall melanophore patterning and subsequent darkening of the skin are similar in *Arthroleptis* and *E. coqui*, there is a difference in the timing of the ventral fusion of the pigmented lateral body walls

This fusion appears much earlier in *Arthroleptis* (TS9, Figure 2) compared to *E. coqui* (TS12, Elinson & Fang, 1998). In *E. coqui* this event correlates with a peak in thyroid gland activity (Laslo et al., 2019) and has experimentally shown to be TH dependent (Callery & Elinson, 2000). In *Arthroleptis*, histologically inferred thyroid activity peaks at TS10/11. Therefore fusion of the lateral body walls seems slightly accelerated. This could be due to a higher TH-sensitivity of the lateral body wall tissue or the earlier expression of TH that cannot be detected histologically. Experimental and molecular/immuno-histochemical or mass spectroscopic data from *Arthroleptis* are needed to clarify this.

### Skin histology in *Arthroleptis* exhibits a phase of metamorphic repatterning that correlates with increased thyroid gland activity

Skin and thyroid gland histology of the *Cardioglossa* tadpole are similar to many other tadpoles at a comparable developmental stage (Cruz & Fabrezi, 2020). Alterations in skin development in *Arthroleptis* therefore seem to be correlated with the evolution of direct development. However, an apical extracellular layer resembling the stratum corneum in adult frog skin is present in the *C. manengouba* tadpole. Tadpoles of various species of *Cardioglossa*, including *C. manengouba*, have been frequently found buried in sediment and other substrate in streams (Blackburn, 2008; Hirschfeld, Barej, Gonwouo, & Rödel, 2012) and this putative stratum corneum could be an adaptation to their fossorial life style.

As reported for *E. coqui* (Fang & Elinson, 1996; Schlosser et al., 1999), major tadpole characters such as cement glands, neuromasts and skein cells (Tamakoshi et al., 1998) were not observed in *Arthroleptis*. Data for *I. henselii* are not available (Goldberg et al., 2020). In *Arthroleptis*, skin development can be divided into four major phases (embryonic, “tadpole”, metamorphic and adult; Figure 8) that are also typical for many biphasic species (Fox, 1986; Robinson & Heintzelman, 1987). At TS4 embryonic ectoderm is a single-cell layered epithelium not yet differentiated into an epidermis. At around TS7 the ectodermal cells have differentiated into a tadpole-like epidermis (two-layered, unicellular glands, inner melanophores). However, the epidermis still exhibits some embryonic features (intracellular yolk granules, low melanophore density, ciliated cells and the lack of a stratum compactum; Figure 8). At TS8, the skin exhibits more mature tadpole-like features such as fewer intracellular yolk granules and ciliated cells, a higher melanophore density, many unicellular glands and a stratum compactum. At TS11, degeneration of the apical epidermal layer and unicellular glands together with the appearance of multicellular gland progenitors and an additional outer melanophore layer (Figure 8) resembles skin repatterning during the metamorphic climax of biphasic species (Fox, 1985; Gaupp, 1904; Verma, 1965). In biphasic species, shortly before metamorphic climax, apical cells in the tadpole epidermis undergo apoptosis except for the single layer of basal cells. The remaining basal cells proliferate extensively during metamorphic climax building up the adult epidermis and gland cells (Kinoshita & Sasaki, 1994; Schreiber & Brown, 2003; Yoshizato, 1992). In *Arthroleptis*, metamorphic skin repatterning is seen at TS11, when the “tadpole” apical epidermal layer degenerates and cells in the remaining basal layer start to proliferate (Figure 8; strong PCNA signal). This patterning of the tadpole-like skin and increased PCNA activity correlate with an increase of histologically inferred thyroid gland activity at TS10/11 (see Figure 6). From TS13 on, the skin of *Arthroleptis* exhibits major features of an adult frog skin, such as a multilayered (three to five layers dorsally) epidermis with an apical stratum corneum, large multicellular glands embedded into a stratum spongiosum and a layer of outer and inner melanophores bordering the thick stratum compactum (Figure 8). This in term correlates with a decreased thyroid gland activity and the absence of a PCNA signal in epidermal cells.

**Figure 8.**
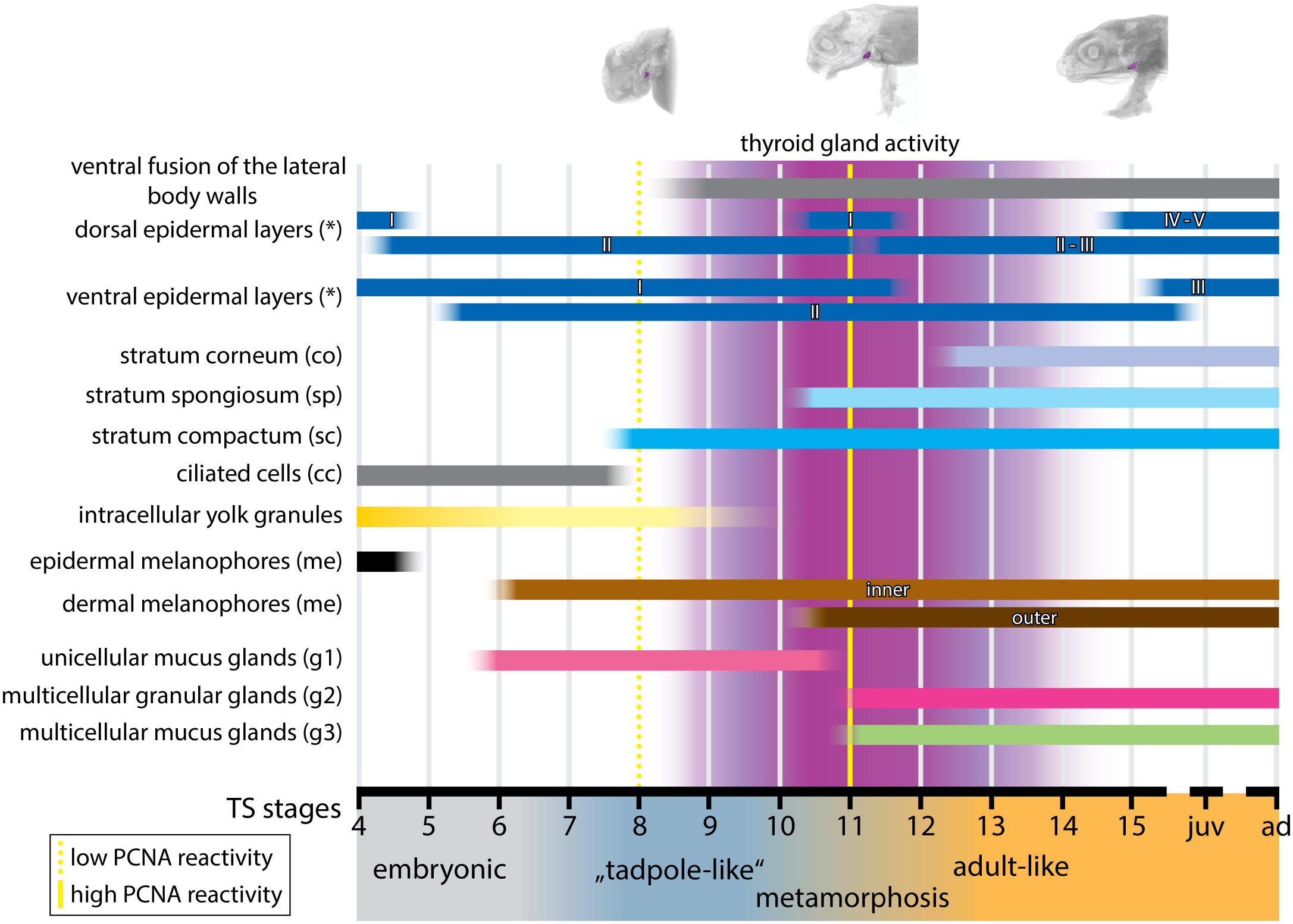
Schematic summary of the skin composition of *Arthroleptis* at different developmental stages. Presumptive thyroid gland activity inferred from histological measurements is indicated in lilac.

Regarding other direct developing species, detailed histological data are only available for two developmental stages of *Ischnocnema henselii* (Goldberg et al., 2020). At TS6, the epidermis of this species is a single-layered epithelium overlaying an undifferentiated mesenchymal dermis (Goldberg et al., 2020). This is very similar to *Arthroleptis* embryos except that the epidermis is two-layered in most body regions at this developmental stage. In *I. henselii* embryos at TS14, epidermal cells start to proliferate forming a two-layered epidermis. Ciliated cells are still present in many body areas, progenitors of multicellular glands are detectable and melanocyte density has increased compared to TS6. The dermis has also differentiated and a stratum compactum is present (Goldberg et al., 2020). This is different to *Arthroleptis*, where the features described for *I. henselii* at TS14 are already present at TS11 (Figure 8). In contrast to *I. henselii*, the short description of a stage likely corresponding to TS9 in *Eleutherodactylus* that exhibits a two-layered epidermis (Adamson, Harrison, & Bayley, 1960), implies an earlier pattern of skin differentiation more similar to *Arthroleptis*. This is very interesting since a differentiated thyroid gland is already present at TS6 in *I. henselii*. This is much earlier compared to *Arthroleptis* (TS8; this study) and *E. coqui* (TS10; Jennings & Hanken, 1998) and skin development does therefore not seem to correlate with thyroid gland maturation in *I. henselii*.

This mismatch between very early TH gland maturation and very late skin remodeling in *I. henselli* could be indicative of a loss of TH-dependency of skin remodeling and hence a loss of constraint. Alternatively, skin maturation might still be dependent on TH-signaling, but TH-production of the thyroid gland could be very low and a concentration necessary to induce skin repatterning is reached very late. Another possibility is that TH-levels are normal but the sensitivity of the skin tissue is decreased, requiring higher TH concentrations to induce repatterning. Both of these TH-dependent scenarios would explain the late onset (i.e. heterochronic shift) of skin remodeling.

### Thyroid gland development and activity in direct developing frogs

As it has been shown that metamorphic changes in anurans, and amphibians in general, are under a complex hormonal control but TH seems to be a major regulator (Shi, 1999). There are two hypotheses how the ancestral thyroid axis has influenced the evolution of direct development (see Jennings & Hanken, 1998 and references therein). (1) Thyroid gland activity and TH production are activated early in development leading to a precocious formation of adult features. Direct development is constrained by the ancestral dependency on TH for adult development. Consequently, direct development evolved by a heterochronic shift (Gould, 1977) of ancestral developmental mechanisms rather than the loss of constraints from it (“heterochronic shift” hypothesis). (2) Metamorphosing tissues lost their ancestral TH-dependency. In this case, the loss of constraints would have been more important in the evolution of direct development than heterochronic shifts of ancestral mechanisms (“loss of constraint” hypothesis).

Experimental studies on *E. coqui* have shown that this species exhibits a mosaic pattern of heterochronic shifts and loss of constraints for different morphological features (Callery & Elinson, 2000; Hanken et al., 1997; Hughes, 1966; Hughes & Reier, 1972; Jennings & Hanken, 1998; Lynn, 1948; Lynn & Peadon, 1955; Townsend & Stewart, 1985).

There are many similarities between *E. coqui* and *Arthroleptis* regarding metamorphic repatterning processes that correlate with increased thyroid activity such as tail regression, remodeling of the “larval” into the adult hyobranchial apparatus and cranial muscle repatterning, (Hanken et al., 1997; Jennings & Hanken, 1998; Schweiger et al., 2017; Schweiger, Naumann, & Müller, in prep.; Townsend & Stewart, 1985). Most of these processes start slightly earlier in *Arthroleptis*, which correlates with an earlier appearance of the thyroid glands compared to *E. coqui* (Hanken & Jennings, 1998; this study). *Ischnocnema henselii* might be another example where some features of skin development might have lost their dependency on TH, but the interpretation of the data for *I. henselii* is hampered by the lack of a detailed investigation of thyroid gland and skin development over several developmental stages.

However, our data indicate that many tadpole specific structures appear, differentiate and are repatterned to the adult configuration during a short metamorphic phase although this cryptic metamorphosis appears to be not as climactic as in biphasic frogs.

### Developmental constraints, heterochrony and the parallel evolution of direct development

Direct development has evolved parallel in different groups of frogs. The comparison of embryonic development of these direct developing frogs offers the possibility to identify how developmental constraints may have influenced the observed pattern of parallel evolution. Many developmental mechanisms are governed by major regulators such as TH. In case of a heterochronic shift of this regulator (e.g. embryonic thyroid activity in direct developing frogs), traits that are under the control of this regulator have to follow this shift. These traits are constrained by this major regulator. Other traits that do not follow the heterochronic shift of this regulator develop normally and are not constrained. This may lead to a decoupling of ancestrally synchronous processes such as, e.g., growth and differentiation of the retinotectal system in *E. coqui* (Schlosser, 2008).

Further investigations of the developmental timing and TH-dependency of anatomical features of *Eleutherodactylus, Arthroleptis, Ischnocnema* and other direct developing frogs are needed. These investigations will clarify if homologous features are generally under the control of TH and therefore constrained by thyroid gland activity. Studies on direct development in frogs, or amphibians in general, and various other cases such as the classic example of the evolution of similar morphotypes in African cichlids (Elmer et al., 2014; Meier, Marques, Wagner, Excoffier, & Seehausen, 2018) or the recently “re-discovered” dispersion/re-aggregation and diapause phases in early killifish development (Furness, Reznick, Springer, & Meredith, 2015; Naumann & Englert, 2018) will help to better understand how developmental processes constrain the evolution of adaptive traits and to further clarify mechanisms and identifying major regulators underlying parallel evolution.

## Experimental Procedures

### Specimens

Embryos of *Arthroleptis wahlbergii* Smith, 1849, collected in South Africa, and *A. xenodactyloides* Hewitt, 1933, collected in Tanzania, were euthanized using 0.1 % tricaine methanesulfonate (MS222, Fluka), fixed in 4% phosphate buffered formalin and stored in 70% ethanol (Schweiger et al., 2017). Staging of embryos follows Townsend and Stewart (1985); TS stages hereafter. One tadpole of *Cardioglossa manengouba* Blackburn, 2008 was available for investigation. For a complete list of specimen see Supplementary material 1.

### Histological sectioning and staining

For sectioning, embryos were dehydrated in an ethanol series (70%, 90%, 2 x 96%; 5 min each), sectioned and subsequently rehydrated (2 x 96%, 90%, 70% ethanol, distilled water; 5 min each) prior to staining. Embryos up to TS8 were embedded in Technovit 8100 (Kulzer, R0010022), sectioned at 3 μm and stained with a mixture of basophilic Methylene blue and acidic Fuchsin (Mulisch & Welsch, 2010). Embryos from TS10/11 on were decalcified for up to three days (Osteomoll, Merck), embedded in Paraplast (Roth, X881.1), sectioned at 8 μm and stained with Heidenhain’s Azan (Mulisch & Welsch, 2010). All sections were made using a Microm HM 360 (Zeiss) and photographs were taken using an Olympus Dotslide BX51 microscope.

### Fluorescent antibody staining

Paraffin sections were rehydrated as described before. For antigen retrieval, slides were transferred to a plastic cuvette with citrate buffer (recipe see Schlosser, 2008) and heated in a microwave (Bosch HMT 72M 420) for 10 minutes at 180W. Subsequently, slides were placed at room temperature (RT, 21-25°C) to cool down for at least 5 minutes and then rinsed in distilled water for 1 minute. Afterwards, slides were rinsed with PBS (3 x 5 minutes), placed into a wet chamber and blocked with antibody diluent (DAKO) for 1 hour at RT. Primary antibodies against proliferating cell nuclear antigen (PCNA-1, 1/200, sc-7907, Santa Cruz; PCNA-2, 1/200, M087901, Dako) were applied overnight at 4°C. On the next day, slides were rinsed in PBS (3 x 5 minutes), blocked and incubated with secondary antibodies (Alexa-488 anti-mouse, #A28175, Thermo Fisher Scientific and Alexa-568 anti-rabbit, #A-11011, Thermo Fisher Scientific) for 1 hour at RT. Afterwards, slides were rinsed with PBS (3 x 5 minutes), quickly washed with distilled water and cover-slipped with Fluoroshield with DAPI (Sigma, F6057). Slides were photographed using a Zeiss Axioplan Microscope equipped with a Spot camera and the Zeiss Axioplan software.

### Micro-CT scanning and 3D reconstruction

Specimens were contrasted using a solution of 1% polymolybdenic acid in 70% ethanol (Metscher, 2009). *Arthroleptis* specimens were CT-scanned using a Nanotom S μCT scanner (Phoenix X-Ray) and the *Cardioglossa* tadpole was studied using synchrotron radiation based x-ray micro-CT. Imaging was performed at the Imaging Beamline P05 (IBL) (Greving et al., 2014; Haibel et al., 2010; Wilde et al., 2016) operated by the Helmholtz-Zentrum-Geesthacht at the storage ring PETRA III (Deutsches Elektronen Synchrotron – DESY, Hamburg, Germany) using a photon energy of 24 keV. Projections were recorded using a CCD camera system (MicroLine ML09000 - Finger Lake Instruments) with an effective pixel size of 2.42 μm. For each tomographic scan 1801 projections at equal intervals between 0 and π have been recorded. Tomographic reconstruction has been done by applying a filtered back projection algorithm (FBP) implemented in a custom reconstruction pipeline (Moosmann et al., 2014) using Matlab (Math-Works) and the Astra Toolbox (Palenstijn, Batenburg, & Sijbers, 2011; van Aarle et al., 2016; van Aarle et al., 2015). For the tomographic reconstruction, raw projections were binned two times resulting in an effective pixel size of the reconstructed volume of 4.83 μm. Three-dimensional reconstructions and mixed surface-volume renderings of the thyroid gland were prepared using AMIRA 5.4.2 (FEI Visualization, Science Group, Bordeaux, France).

### Follicle cell height and follicle diameter measurements

Qualitative and quantitative descriptions of thyroid glands are based on sections from the middle region of the paired thyroid lobes according to Cruz and Fabrezi (2020). Measurements of follicle cell heights and follicle diameter were obtained from digitalized sections using ImageJ (Schneider, Rasband, & Eliceiri, 2012). A box and whisker plot of follicle cell heights was prepared using Microsoft Excel 2010. Numbers of follicles were counted and mean values of the measured follicle diameters were calculated for each specimen investigated.

### Photographs and Image processing

Photographs of whole embryos were taken using a Zeiss Discovery V12 stereomicroscope with an attached Zeiss AxioCam digital camera. Brightness and Contrast of Images was adjusted using either ImageJ or Adobe Photoshop CS6. Channel colors of fluorescent antibody staining were assigned in ImageJ. In some images the “CLAHE” filter implemented in ImageJ was applied to enhance local contrast.

## Acknowledgements

We would like to thank Katja Felbel for valuable help in the laboratory. We thank Mark-Oliver Rödel for critical reading of the manuscript and a photograph of an adult *C. manengouba* and Mareike Hirschfeld for a photograph of a *C. manengouba* tadpole. We thank Lennart Olsson for critical reading of the manuscript. We thank Paul Lukas for help with the Dot Slide Microscope. We would also like to thank Christoph Englert, Birgit Perner and Dagmar Kruspe for the kind gift of the two PCNA antibodies.

---

**Supplementary material 1.**
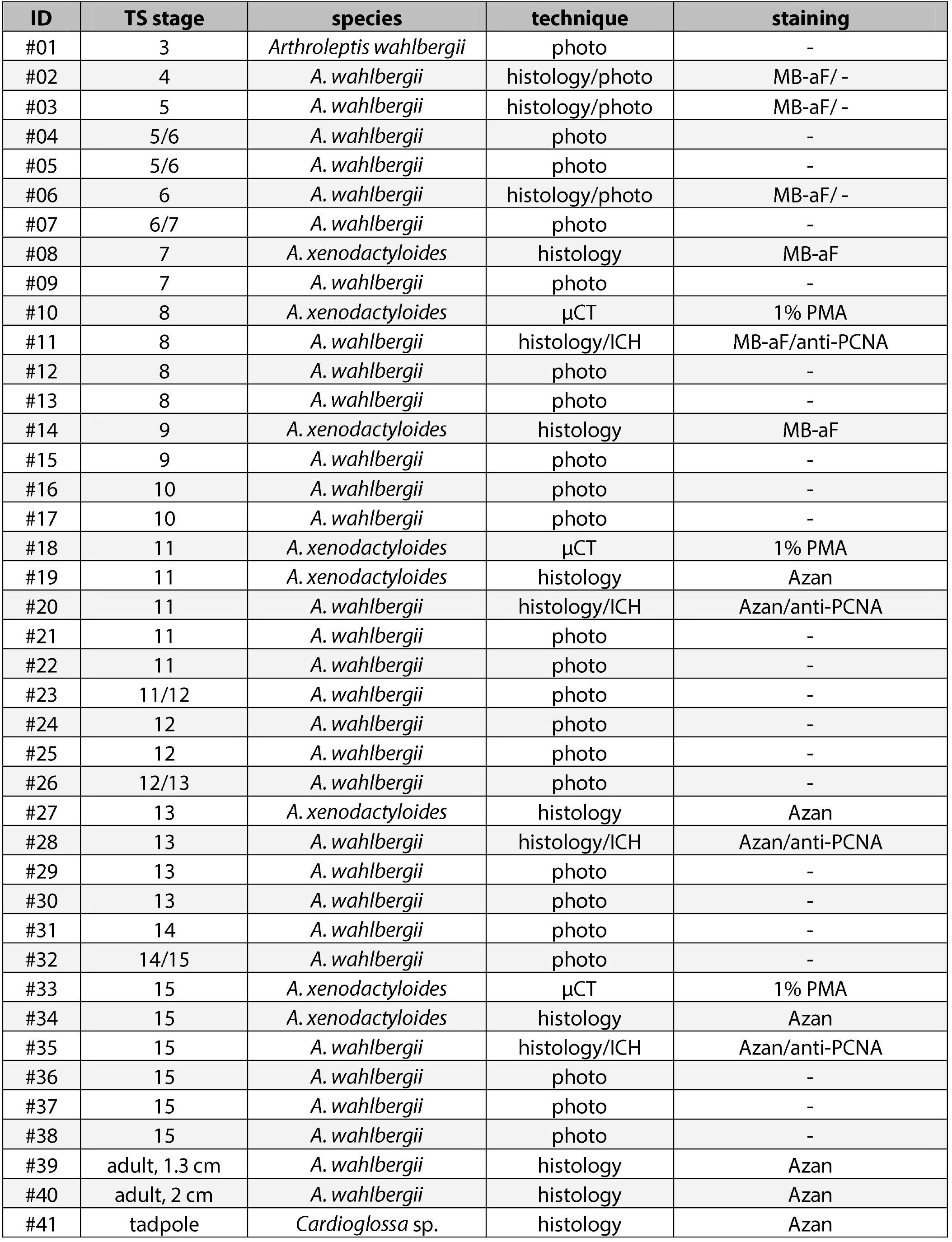
Specimen list. ICH, immune-histochemistry; MB-aF, Methylene blue acidic Fuchsin; PMC, polymolybdenic acid; μCT, micro-computed tomography.

**Supplementary material 2.**
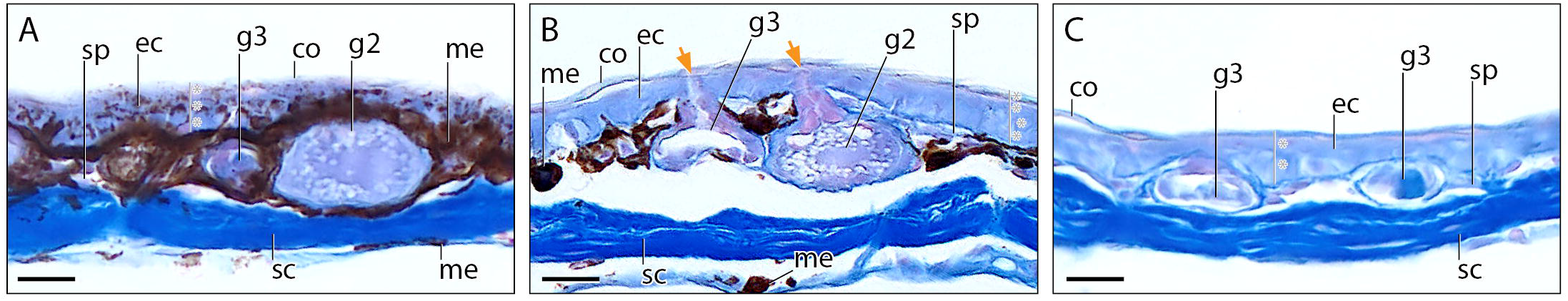
Adult skin histology. Histological cross sections of the skin of an adult (1.3 cm) *Arthroleptis wahlbergi*. A, dorsal head region. B, lateral head region. The orange arrow indicates the glandular duct. C, ventral head region. The number of epidermal cellular layers is indicated by a grey line and asterisks. Scale bar is 20 μm. co, stratum corneum; ec, epithel cell; g2, multicellular granular gland; g3, multicellular mucus gland; me, melanophore; sc, stratum compactum; sp, stratum spongiosum.

